# Giant pyramidal neurons of the primary motor cortex express vasoactive intestinal polypeptide (VIP), a known marker of cortical interneurons

**DOI:** 10.1101/2024.01.22.576584

**Authors:** Sadaf Teymornejad, Katrina H. Worthy, Marcello G. P. Rosa, Nafiseh Atapour

## Abstract

Vasoactive intestinal polypeptide (VIP) is known to be present in a subclass of cortical interneurons that are involved in the disinhibition of excitatory pyramidal neurons. Here, using three different antibodies, we demonstrate that VIP is also present in the giant layer 5 pyramidal (Betz) neurons which are characteristic of the limb and axial representations of the marmoset primary motor cortex (cytoarchitectural area 4ab). No VIP staining was observed in smaller layer 5 pyramidal cells present in the primary motor facial representation (cytoarchitectural area 4c), or in premotor cortex (e.g. the caudal subdivision of the dorsal premotor cortex, A6DC), indicating the selective expression of VIP in Betz cells. The most intense VIP staining occurred in the largest Betz cells, located in the medial part of A4ab. VIP in Betz cells was colocalized with neuronal specific marker (NeuN) and a calcium-binding protein parvalbumin (PV). PV also intensely labelled axon terminals surrounding Betz cell somata. Whereas Betz cells bodies were located in layer 5, VIP-positive (VIP^+^) interneurons were more abundant in the superficial cortical layers. They constituted about 5-7% of total cortical neuronal population, with the highest density observed in area 4c. Our results demonstrate the expression of VIP in the largest excitatory neurons of the primate cortex, which may offer new functional insights into the role of VIP in the brain, and also provide opportunities for genetic manipulation of Betz cells.

## 1. INTRODUCTION

Similar to other primates (e.g. Vogt and Vogt, 1919; Matelli et al., 1985; Watanabe-Sawaguchi et al., 1991) in the marmoset monkey the primary motor cortex (M1, Brodmann Area 4) is formed by distinct cytoarchitectural subdivisions. According to the contemporary view the representations of the limb and body axial musculatures are encompassed within cytoarchitectural area 4ab, located medially, whereas the representations of the head musculature (including facial and oral) are located laterally, in cytoarchitectural area 4c (Burman et al., 2008: Paxinos et al., 2012). In particular, A4ab contains the unique “gigantopyramidal” neurons, known as Betz cells in primates (Bakken et al., 2021; Betz, 1874; Jacobs et al., 2018). In contrast, in A4c layer 5 is populated by smaller pyramidal neurons (Burman et al., 2008). Betz cells are characterised by their pyramidal shape, large size, and prominent apical dendrites that are oriented along a vertical axis. Although more sparsely distributed, they can be as 20 times larger in volume when compared to other pyramidal cells (in humans; Rivara et al., 2003).

Betz cells have specialised molecular and physiological characteristics (Bakken et al., 2021). Given their distinctive action-potential properties (Spain et al.,1991; Vigneswaran et al, 2011) and direct connectivity onto the ventral horn of the spinal cord (Lemon, 2008), they are well placed for initiation and modulation of fine movement. Betz cells can be evidenced using markers of pyramidal neurons, such as antibodies against non-phosphorylated neurofilament (e.g. SMI-32; Tsang et al., 2000; Szocsics et al., 2021), as well as general markers of neuronal nuclei (e.g. NeuN; Szocsics et al., 2021). In addition, some of these cells express specifically nitric oxide synthase (NOS, Wallace et al., 1996) and/ or the calcium binding protein parvalbumin (PV, Szocsics et al., 2021).

There is a large body of evidence to indicate that the local vasoactive intestinal polypeptide-positive (VIP^+^) interneurons exert inhibition onto excitatory pyramidal cells, either directly or indirectly, to shape their outputs (Lee et al., 2013; Garcia-Junco-Clemente et al. 2017; Pi et al., 2013: Zhou et al., 2017; Yetman et al. 2019; Szadai et al, 2022). Here, we present evidence that staining of M1 with antibodies against VIP reveals not only a subset of GABAergic interneurons, as expected from previous studies (Gabbott and Bacon 1997; Peters et al., 1987; Prönneke et al 2015; Kim et al.., 2017), but also Betz cells. The spatial and laminar distribution of VIP^+^ interneurons and Betz cells are compared.

## 2. MATERIALS AND METHODES

### 2.1 Animals

Materials were sourced from nine healthy adult marmosets (*Callithrix. jacchus*), as detailed in Table 1. The experiments were conducted in accordance with the Australian Code of Practice for the Care and Use of Animals for Scientific Purposes. All procedures were overseen by the Animal Ethics Experimentation Committee at Monash University. The animals had no veterinary record of serious acute, or chronic health conditions. Some of the animals were also used anatomical tracing experiment unrelated to the present study (Majka et al., 2016, 2020; for details of the tracer experiments, see www.marmosetbrain.org).

**Table 1.** Details of sex, age, and staining.

| Subject (Sex) | Neuronal marker | Staining method | Age at perfusion (months) |
| --- | --- | --- | --- |
| CJ195 (M) | PV | IHC | 43 |
| CJ200 (F) | PV, CB, CR, NeuN | IHC | 43 |
| CJ205 (M) | CB, CR | IHC | 43 |
| CJ226 (F) | VIP, NeuN | IHC/IF | 24 |
| CJ227 (M) | VIP, NeuN | IHC/IF | 37 |
| CJ232 (F) | VIP, PV | IF | 30 |
| CJ240 (M) | VIP, NeuN | IHC/IF | 36 |
| CJ249 (F) | VIP, GABA | IF | 36 |
| F1741 (F) | CB, CR, NeuN | IHC/IF | 40 |

*Abbreviations:* PV, Parvalbumin; CB, Calbindin; CR, Calretinin; VIP, Vasoactive intestinal polypeptide; NeuN, Neuronal nuclei; GABA, Gamma-aminobutyric acid; IHC, Immunohistochemistry using DAB; IF, Immunofluorescence.

### 2.2 Tissue perfusion and processing

The marmosets were euthanized with an overdose of sodium pentobarbitone (100 mg/kg, i.m.) and transcardially perfused with 0.1 M heparinized phosphate buffer (PBS; pH 7.2), followed by 4% paraformaldehyde in 0.1 PBS. The brains were dissected and post-fixed in the same medium for one hour prior to immersion in phosphate buffered saline, with increasing concentrations of sucrose (10%, 20% and 30%), over several days. Using a cryostat, frozen 40μm coronal sections were obtained. In cases also used for anatomical tracing, the sections used for the present analyses were sourced from the hemisphere contralateral to the tracer injections. Five sequential series were collected, with the first two being stained for myelin (Gallyas, 1979; modified by Worthy et al., 2017), and for cell bodies (cresyl violet Nissl stain). These series were used to delineate boundaries of motor cortex (Burman et al. 2008). The other series were used for immunostaining, as detailed in Table 1.

Immunostainings were conducted by incubating sections in blocking solution (10% normal horse serum and 0.3% Triton-X100 in 0.1 M PBS) for 1 hr at room temperature before undergoing incubation in primary antibody at 4°C for 42–46 h (Table 2). For immunohistochemistry using DAB (IHC), a biotinylated horse anti-mouse IgG secondary antibody incubation was applied for 30 min, followed by treatment with Avidin-Biotin Complex (ABC) reagent and DAB (3,3’-Diaminobenzidine) substrate working solution. Alexa Fluor secondary antibodies were used for immunofluorescence staining (IF) (Table 2).

**Table 2.**
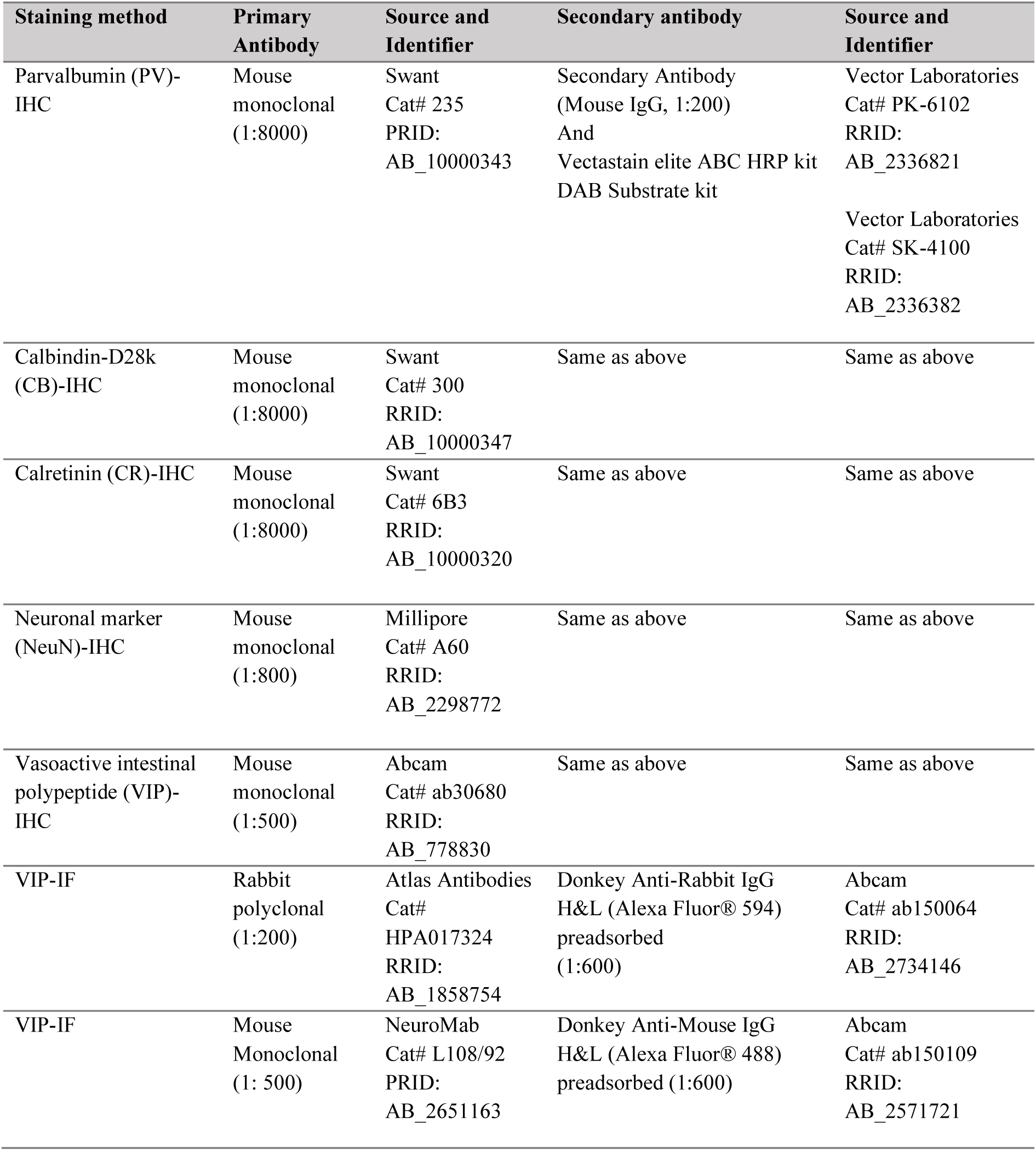
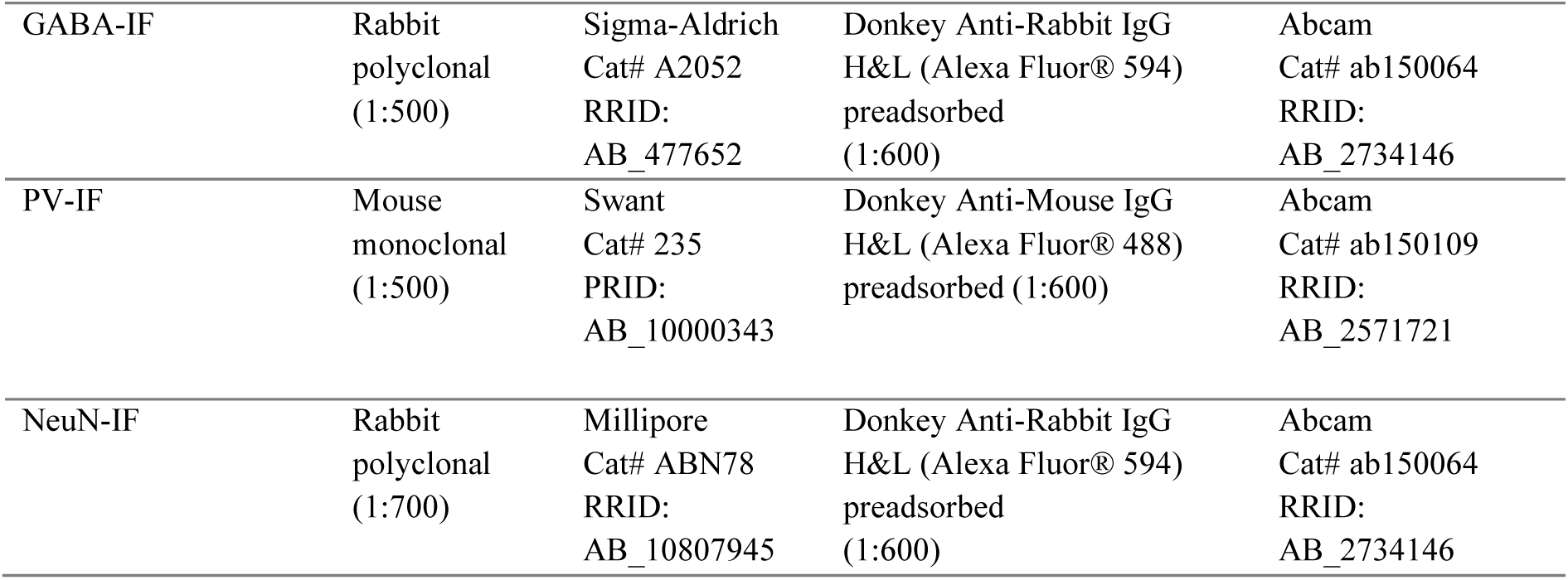
List of primary and secondary antibodies used for immunostaining.

| Staining method | Primary Antibody | Source and Identifier | Secondary antibody | Source and Identifier |
| --- | --- | --- | --- | --- |
| Parvalbumin (PV)-IHC | Mouse monoclonal (1:8000) | Swant<br>Cat# 235<br>PRID:<br>AB_10000343 | Secondary Antibody (Mouse IgG, 1:200)<br>And<br>Vectastain elite ABC HRP kit<br>DAB Substrate kit | Vector Laboratories<br>Cat# PK-6102<br>RRID:<br>AB_2336821<br>Vector Laboratories<br>Cat# SK-4100<br>RRID:<br>AB_2336382 |
| Calbindin-D28k (CB)-IHC | Mouse monoclonal (1:8000) | Swant<br>Cat# 300<br>RRID:<br>AB_10000347 | Same as above | Same as above |
| Calretinin (CR)-IHC | Mouse monoclonal (1:8000) | Swant<br>Cat# 6B3<br>RRID:<br>AB_10000320 | Same as above | Same as above |
| Neuronal marker (NeuN)-IHC | Mouse monoclonal (1:800) | Millipore<br>Cat# A60<br>RRID:<br>AB_2298772 | Same as above | Same as above |
| Vasoactive intestinal polypeptide (VIP)-IHC | Mouse monoclonal (1:500) | Abcam<br>Cat# ab30680<br>RRID:<br>AB_778830 | Same as above | Same as above |
| VIP-IF | Rabbit polyclonal (1:200) | Atlas Antibodies<br>Cat#<br>HPA017324<br>RRID:<br>AB_1858754 | Donkey Anti-Rabbit IgG H&L (Alexa Fluor® 594) preadsorbed (1:600) | Abcam<br>Cat# ab150064<br>RRID:<br>AB_2734146 |
| VIP-IF | Mouse Monoclonal (1: 500) | NeuroMab<br>Cat# L108/92<br>PRID:<br>AB_2651163 | Donkey Anti-Mouse IgG H&L (Alexa Fluor® 488) preadsorbed (1:600) | Abcam<br>Cat# ab150109<br>RRID:<br>AB_2571721 |
| GABA-IF | Rabbit polyclonal (1:500) | Sigma-Aldrich Cat# A2052<br>RRID: AB_477652 | Donkey Anti-Rabbit IgG H&L (Alexa Fluor® 594) preadsorbed (1:600) | Abcam Cat# ab150064<br>RRID: AB_2734146 |
| PV-IF | Mouse monoclonal (1:500) | Swant Cat# 235<br>PRID: AB_10000343 | Donkey Anti-Mouse IgG H&L (Alexa Fluor® 488) preadsorbed (1:600) | Abcam Cat# ab150109<br>RRID: AB_2571721 |
| NeuN-IF | Rabbit polyclonal (1:700) | Millipore Cat# ABN78<br>RRID: AB_10807945 | Donkey Anti-Rabbit IgG H&L (Alexa Fluor® 594) preadsorbed (1:600) | Abcam Cat# ab150064<br>RRID: AB_2734146 |

Subsequently, Aperio Scanscope AT Turbo (Leica Biosystems) was used to scan myelin, Nissl and IHC stained sections. Fluorescent Images from the motor cortex were captured using a Nikon C1 laser scanning confocal microscope. A comprehensive list of all antibodies used in this experiment, along with their respective dilution rates, is presented in Table 2.

### 2.3 Quantification of VI**P^+^** interneurons and statistical analysis

The three distinct regions of the primary motor cortex, namely areas 4a, 4b (encompassing representations of the axial and limb musculatures) and 4c (the representation of the head musculature), along with the caudal subdivisions of the dorsal premotor area (A6DC), were delineated using the criteria defined by Burman et al. (2008; 2014a, b).

Four VIP-stained sections were selected from each subdivision of the primary motor cortex from three cases that underwent IHC (CJ226, CJ227 and CJ240). Using standard features of the Aperio Image Scope software, square counting frames (150μm × 150μm) were aligned across the cortical thickness. The uppermost frame was aligned between the border of layers 1 and 2. The frames were equidistant (10μm apart) and covered all cortical layers of the cortex up to the white matter (Atapour et al. 2019), the number of counting frames varied depending on the thickness of the cortex in each section. Each counting frame was bounded by 2 exclusion lines (left and bottom) and 2 inclusion lines (right and top), as previously described (Atapour et al., 2019). A cell was counted if it was completely within the counting frame or if the cell body contacted the inclusion lines.

The number of counted cells per frame was averaged in each section and used to calculate the neuronal density. The section thickness (40μm) and a shrinkage factor of 0.801 obtained in a previous study (Atapour et al., 2017) were considered when calculating the number of cells per volume unit. For the analysis presented in Fig. 5**e**, we measured the VIP fluorescence intensity of several neurons in one section as a relative quantity. We further calculated the corrected total fluorescence [CTF = integrated density - (area of selected cell × mean fluorescence background, Fig 5**f**)] to best access the amount of VIP expression in the two cell types.

Statistical analysis involved the use of unpaired t test or one-way ANOVA followed by post hoc Tukey’s tests. Statistically significant differences were those with a p-value < 0.05.

## 3. RESULTS

Immunostaining against VIP in the motor cortex led to the visualisation of giant Betz cells of layer 5, as well as a subset of putative inhibitory neurons (Fig 1). While the expression of VIP in interneurons has been demonstrated in other species (Gabbott and Bacon 1997; Peters et al., 1987; Prönneke et al 2015; Szadai et al, 2022; Tremblay et al., 2016), our data also indicate VIP expression in Betz cells, which was confirmed using three different antibodies (Table 2). The VIP antibody used in different analyses is given in the corresponding figure legends.

**Figure 1.**
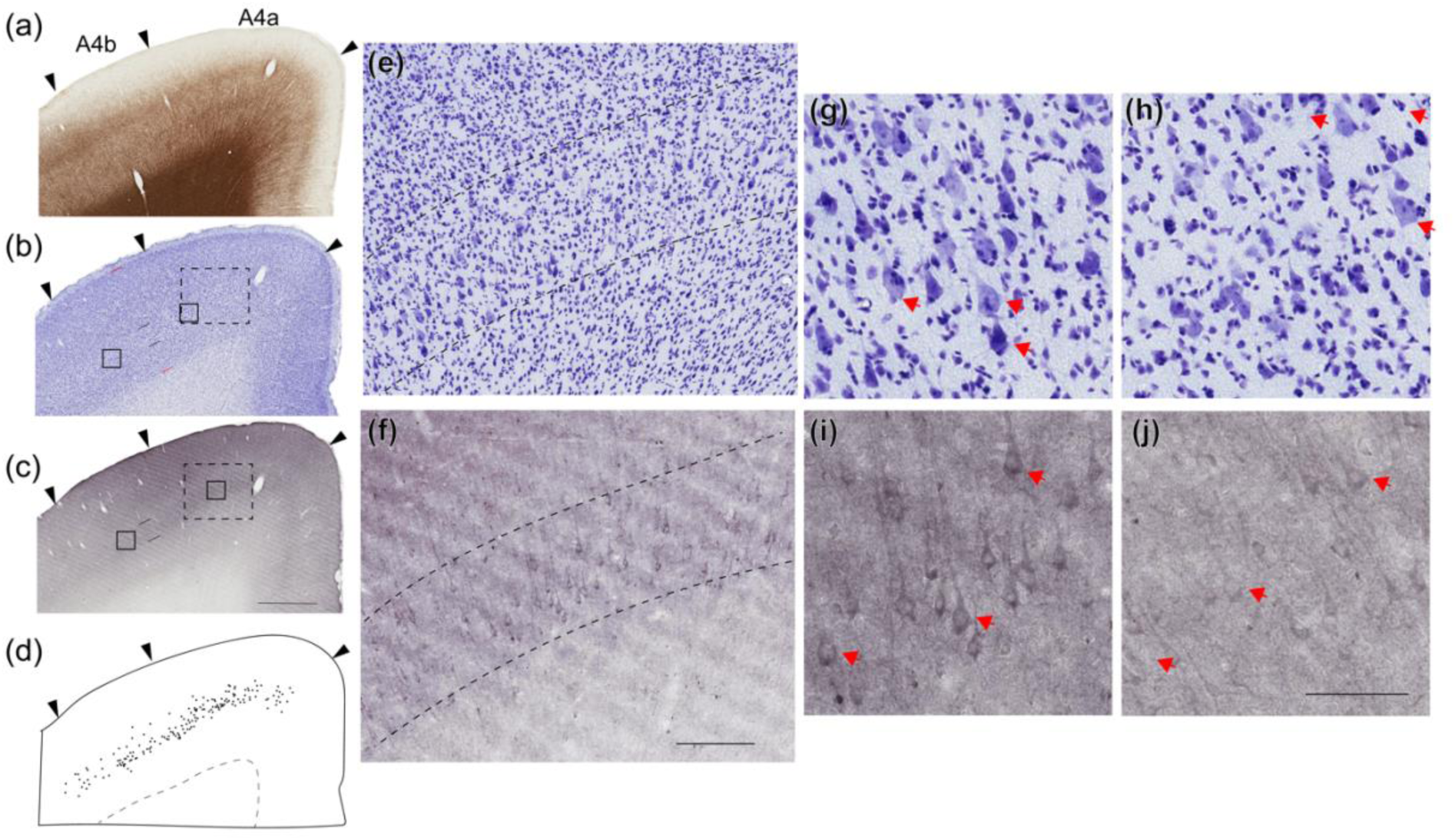
Vasoactive intestinal polypeptide (VIP) is expressed in giant pyramidal (Betz) cells of primary motor cortex. Representative images of primary motor cortex (M1, Area 4a and Area 4b) are shown using myelin (a), nissl (b) and VIP staining (c) in CJ226. Arrowheads point to the areal boundaries (Paxions et al., 2012). Red lines in b & c identify layers 2 to 6 while layer 5 is indicated by small black lines. Schematic of stained layer 5 Betz cells is indicated in d. Betz cells are clearly visible within layer 5 (between the two dashed lines) in e & f representing the area within the dashed rectangles in b & c. Red arrows point to Betz neurons from A4a (g & i) and A4b (h & j) as marked in b & c. Scale: 1 mm for a-d, 200µm for e-f & 100µm for g-j. VIP antibody used in production of images is from Abcam (Cat# ab30680).

### 3.1. Betz cells

VIP staining intensity was strongest in the largest layer 5 pyramidal cells of A4ab, which are located more medially in this area, as defined using Nissl and myelin staining (Fig 1**a-c**, Paxinos et al., 2012). This region corresponds to the representations of the leg and trunk musculatures (Burman et al., 2008). The locations of stained Betz cells are shown by black circles in Fig 1**d**. VIP staining intensity in Betz cells diminishes toward the more lateral parts of A4b (Fig 1**e-j**). The cell body and apical dendrites of Betz cells were clearly visible in the VIP staining, similar to the observations with Nissl staining, yet with more pronounced delineation of the apical dendrite (Fig 1**e-j**). There was no VIP staining in layer 5 pyramidal cells of A4c and premotor area 6DC (Fig 2), which lack the gygantopyramidal characteristics (Burman et al. 2008, 2014).

**Figure 2.**
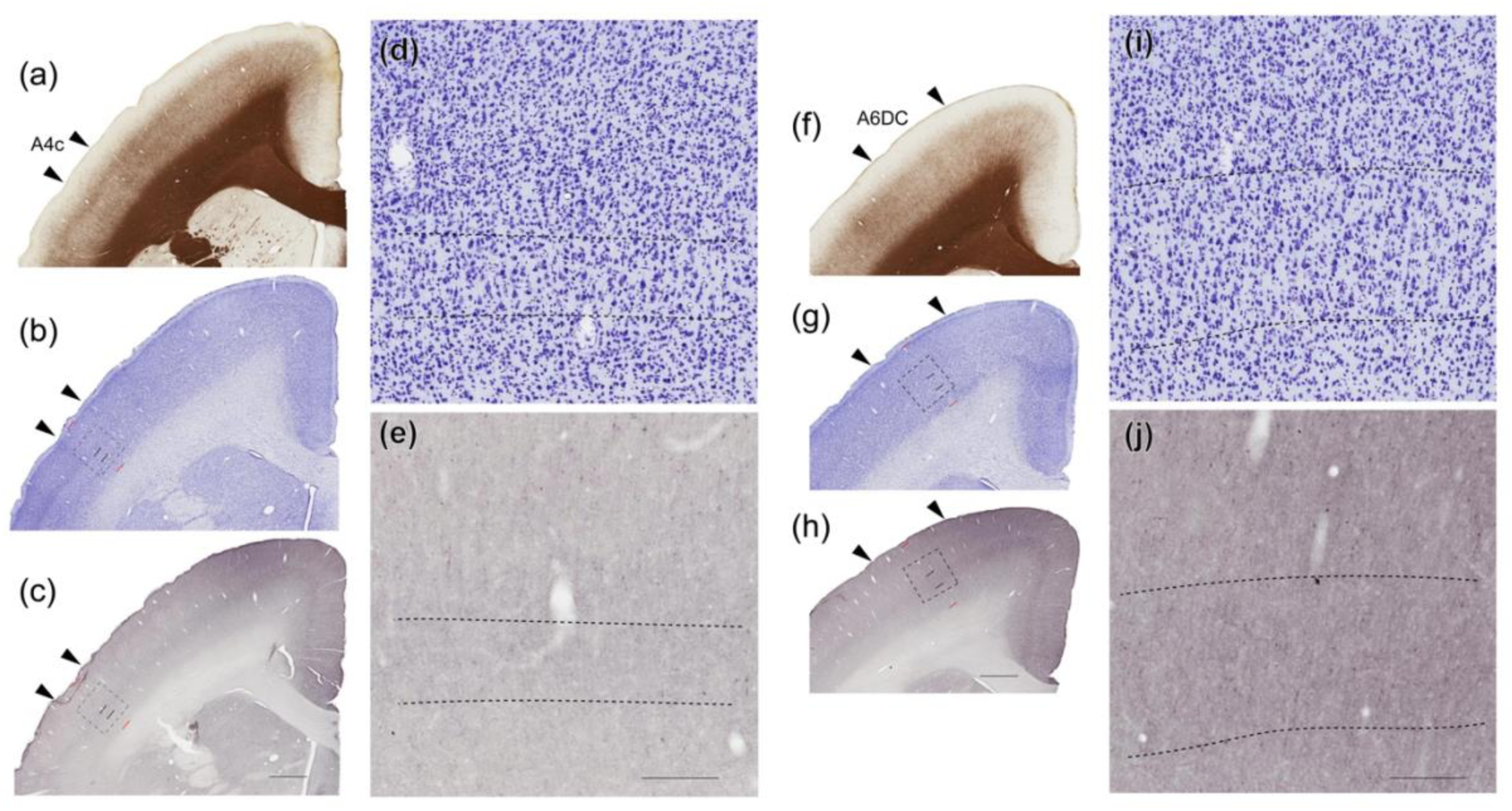
Expression of vasoactive intestinal polypeptide (VIP) is limited to the largest layer 5 pyramidal cells. Representative images of Area 4c (a-e) and A6DC (caudal subdivisions of the dorsal premotor area, f-j) are shown using myelin (a & f), nissl (b & g) and VIP staining (c & h) in CJ226. Arrowheads point to the areal boundaries (Paxions et al., 2012). Red lines in b-c & g-h identify layers 2 to 6, while layer 5 is indicated by small black lines. Large pyramidal cells are only visible within layer 5 (between the two dashed lines) in Nissl staining (d & i; representing area within the dashed rectangle in b & g, respectively). Scale: 1 mm for a-c & f-h, 200µm for d-e & i-j. VIP antibody used in production of images is from Abcam (Cat# ab30680).

We confirmed the presence of VIP staining for the Betz cells by colocalization with the neuronal marker (NeuN), and PV (Fig 3 & 4). PV-positive (PV^+^) axonal endings, which are known to heavily innervate Betz cells (Szocsics et al., 2021), were very evident in our material, forming perisomal networks that delineated the Betz cell bodies (Fig 3). We also found that PV is present in the soma of a subset of Betz cells in marmoset, as reported previously in human brain (Szocsics et al., 2021). The PV signal was far less intense in Betz cell bodies, compared to interneurons or PV^+^ terminal nets delineating soma boundaries, but was clearly distinguishable from the background, and particularly evident by comparison with PV negative (PV^-^) Betz cells. Examples of PV^+^ and PV^-^Betz cells alongside interneurons, are shown in Fig 4. The two other calcium binding proteins tested, calbindin D28-K (CB) and calretinin (CR) were not expressed within Betz cells (Fig 4). This is despite the known expression of CB in subpopulations of pyramidal cells in primates, including in the marmoset (Kondo et al., 1999, Atapour et al., 2024). The largest of Betz cells reached up to 25µm in soma diameter in the marmoset brain (Fig 3 & 4).

**Figure 3.**
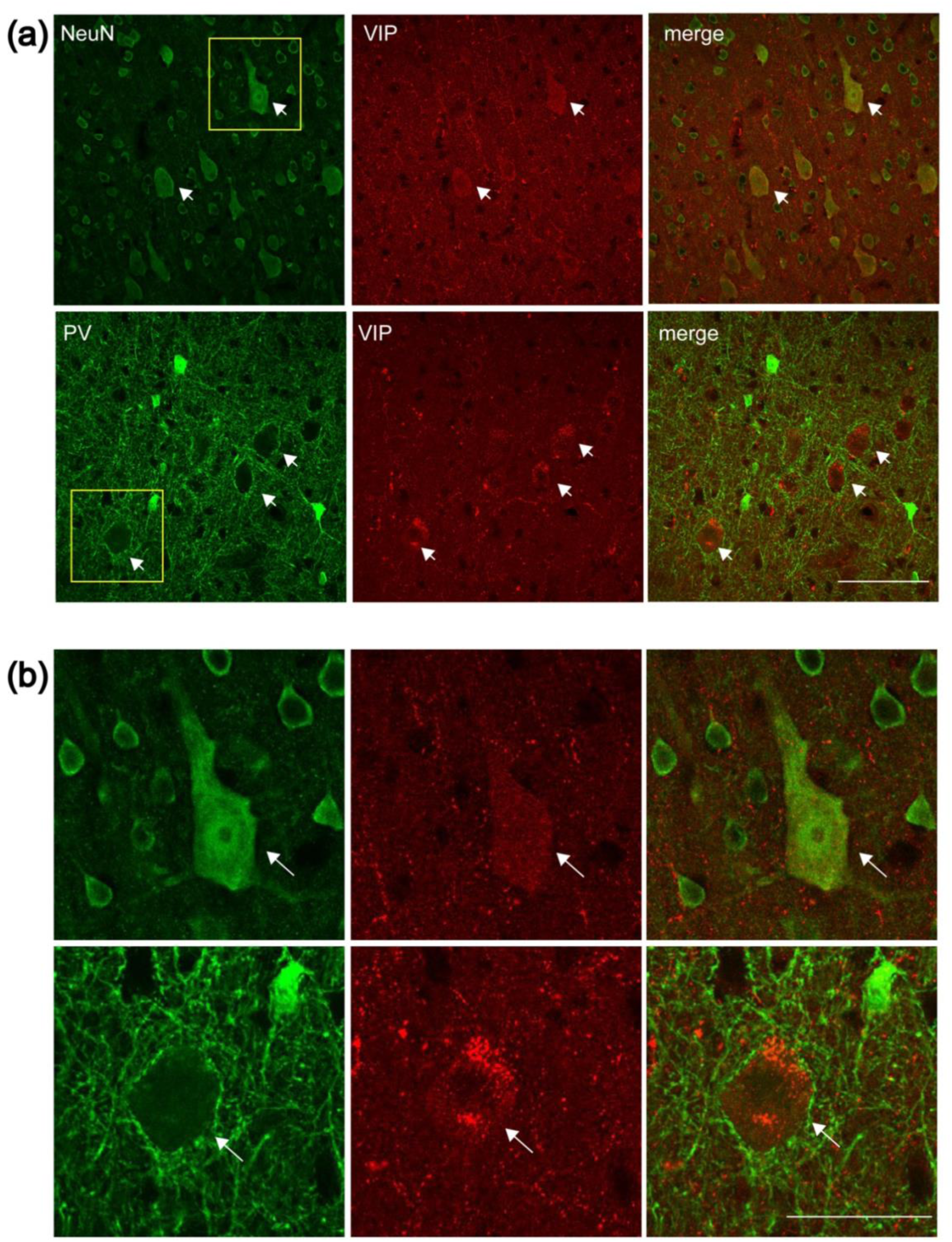
Co-localisation of vasoactive intestinal polypeptide (VIP) with neuronal marker (NeuN) and parvalbumin (PV) in the Betz cells. (a) Confocal images of cortical layer 5 showing immunoreactivity to VIP either with neuronal marker NeuN (top) or PV (bottom). (b) Insets at higher magnification. White arrows point to the Betz cells in all images. Scale: 100µm (a) and 50µm (b). VIP antibody used in production of images is from Atlas Antibodies (Cat# HPA017324).

**Figure 4.**
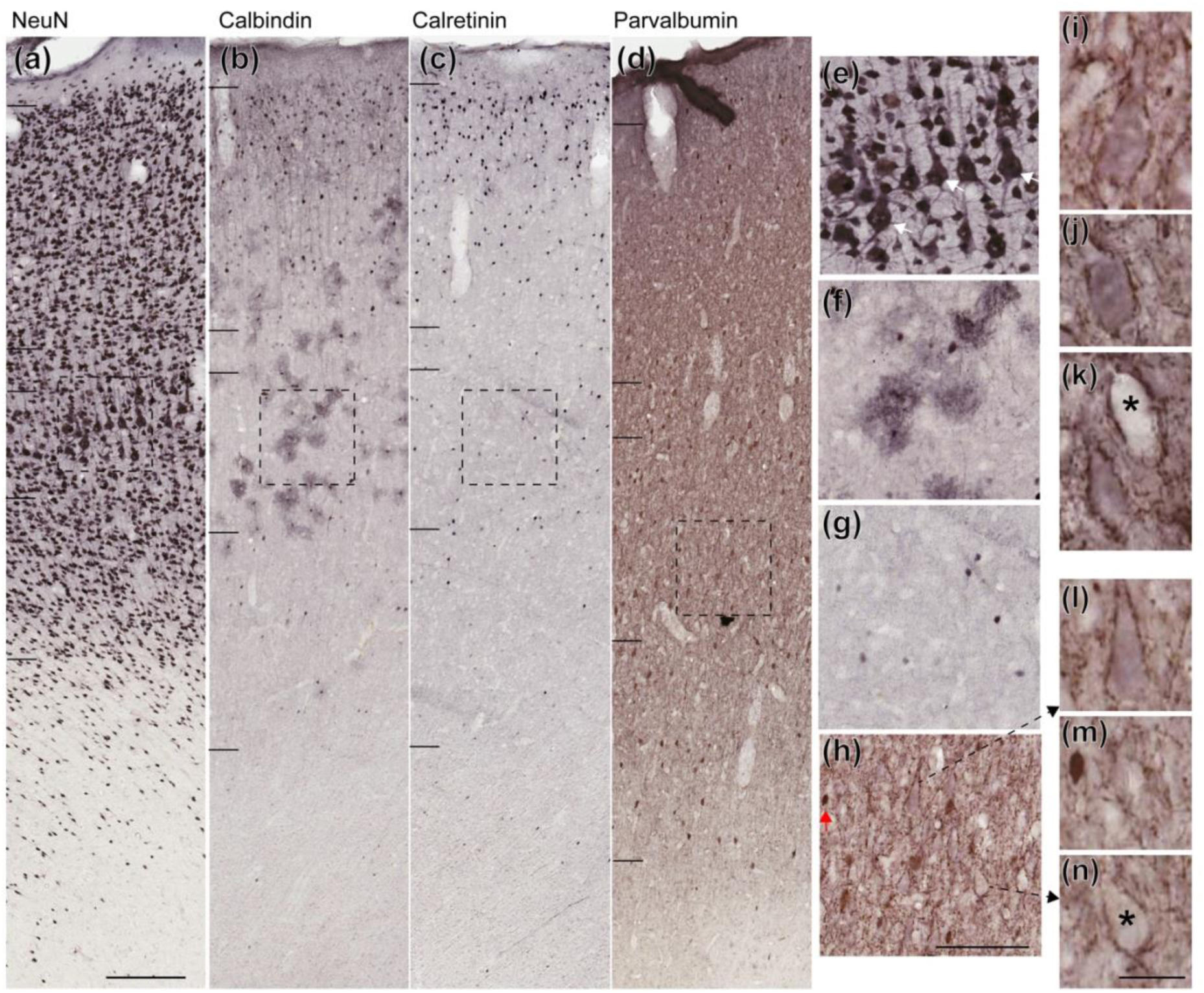
Giant pyramidal cells of the primary motor cortex are identified by parvalbumin (PV) but no other calcium binding proteins. Examples of NeuN (a), calbindin (b), calretinin (c) and parvalbumin (d) staining of area 4A. The boxed areas (dashed squares) in a-d are shown in higher magnification (e-h), respectively. White arrows in e point to Betz cells. Betz cells in i-k are taken from sections not shown here and those in l & n are magnified from panel h. Asterisks (in panels k & n) point to Betz cells that do not contain PV in their soma. PV content is also compared to an interneuron visible in m, highlighted by red arrow in panel h. Scale: 200µm for a-d, 100µm for e-h & 25µm for i-n.

### 3.2. VIP+ inhibitory neurons

VIP staining revealed a subset of inhibitory neurons in the marmoset motor cortex (Fig 5**a-c**). VIP^+^ interneurons had the morphology of dendritic targeting cells, likely including bipolar cells, double bouquet cells and bitufted cells (Markram 2004, Fig 5**a**). Co-staining for VIP and GABA confirmed the inhibitory nature of VIP^+^ interneurons (Fig 5**b-c**). The arbitrary fluorescence intensity was much higher in the VIP^+^ interneurons than in VIP^+^ Betz cells (Fig 5**d-e**, unpaired t-test; 3165 ± 281 vs. 1333 ± 125, p<0.0001). However, considering somatic area, the corrected total fluorescence within the cell bodies was similar (Fig 5**f**, unpaired t test, 18347 ± 2473 vs. 14404 ± 258, p = 0.16).

**Figure 5.**
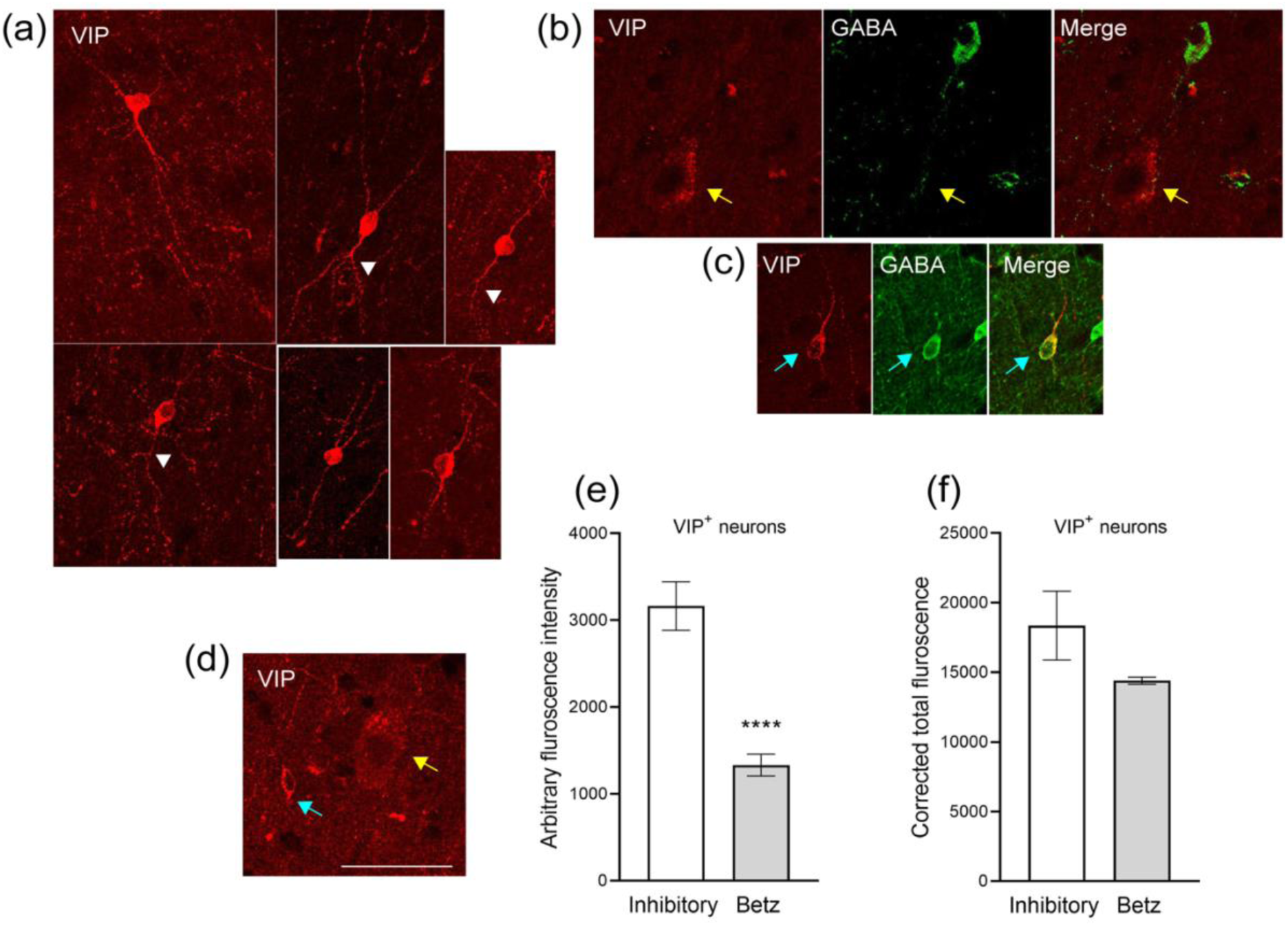
Antibody against vasoactive intestinal polypeptide (VIP) stains inhibitory interneurons as well as Betz cells. Representative images of different morphologies of VIP-positive (VIP^+^) inhibitory neurons of the primary motor cortex (a). White arrowheads point to the dendritic bifurcations. Inhibitory nature of VIP^+^ interneurons is shown by co-localisation with gamma aminobutyric acid (GABA) that is missing in Betz cells (b-c). Fluorescence intensity comparison for a VIP^+^ inhibitory neuron and Betz cell (d). Cyan and yellow arrows point to VIP^+^ inhibitory neuron and Betz cell respectively in b-d. Mean ± SEM density of arbitrary fluorescence intensity obtained from variable somatic locations (e) and corrected total fluorescence for somatic area (f) for VIP^+^ inhibitory and Betz neurons. Scale: 50µm. VIP antibodies used in production of images are from Atlas Antibodies (Cat# HPA017324, a & d) and Neuro Mab (Cat# L108/92, b-c).

In all subfields of primary motor cortex, VIP^+^ inhibitory neurons were mostly concentrated in the supragranular layers, and their density gradually decreased toward layer 6 (Fig 6**a-c**). The mean average density of VIP^+^ interneurons was significantly higher in the A4c compared to the other subdivisions of primary motor cortex [Fig 6**d**, one-way ANOVA: F (2, 33) = 21.26, p< 0.0001; Tukey’s test; A4c (4745±146/mm^3^) vs. A4a (3065± 224/mm^3^) or A4b (3410± 199/mm^3^), p< 0.0001)]. VIP^+^ interneurons corresponded to 4.5, 5.2 and 6.6% of the neuronal population in cytoarchitectural subdivisions A4a, A4b and A4c, respectively (Fig 6**e**).

**Fig 6.**
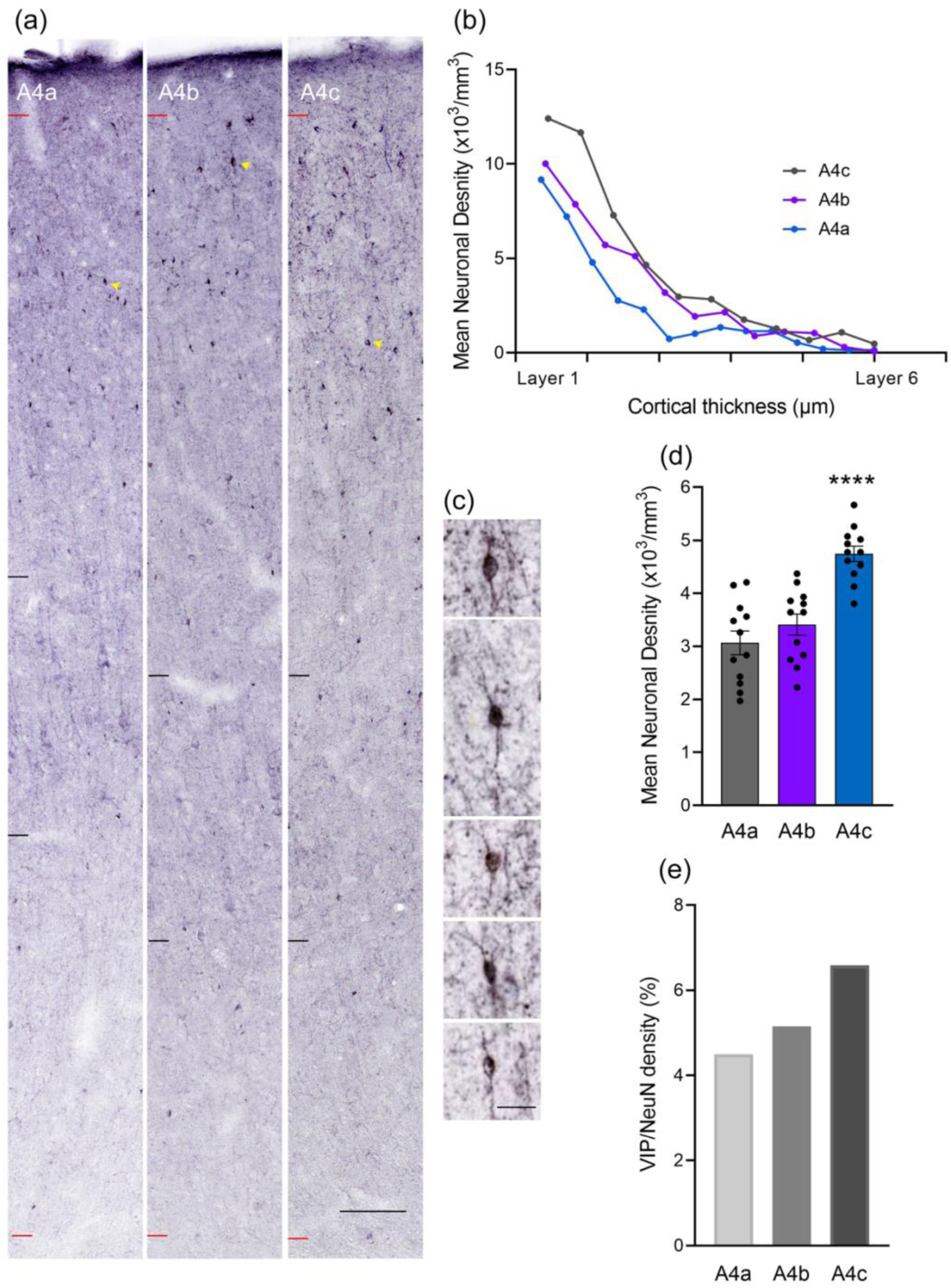
Layer profile of vasoactive intestinal polypeptide-positive (VIP^+^) interneurons of the primary motor cortex. Representative VIP-stained strips from different parts of primary motor cortex including areas A4a, A4b and A4c (a). Small red lines identify the border with layer 2 and white matter. Limits of layer 5 are shown by small black lines. Quantification of VIP^+^ interneuron density across cortical depth (b). Individual cells are indicated by yellow arrows. Some VIP^+^ interneurons are shown at higher magnifications in c. Scale: 100µm in and 20µm in c. Mean ± SEM density of VIP^+^ interneurons (d) and its percentage (e, as ratio of total neurons obtained by NeuN staining in our previous study, Atapour et al., 2019). Black circles represent individual sections in d. VIP antibody used in production of images is from Abcam (Cat# ab30680).

## 4. DISCUSSION

The primary motor cortex is not only involved in motor control, but also in sensory integration, behavioural strategizing, working memory, and decision-making (Ebbesen et al., 2018). Its susceptibility in several disease including neurodegenerative conditions and ageing (Cleveland et al., 2001; Hof & Perl 2002; Scheibel et al., 1977; Seeley 2008; Udaka et al., 1986) warrant further understanding of its neuronal neurochemistry. Here we demonstrate that VIP, a peptide with widespread distribution in many organs, is expressed in the largest excitatory pyramidal neurons of the primate neocortex, adding to the intricacy of distinctive gene-expression and physiological features of the giant layer 5 Betz cells (Bakken et al., 2021). This finding extends VIP expression to a new class of cortical neurons, raising new questions about the physiological role of this peptide in the motor cortex.

### 4.1. VIP expression in Betz cells

VIP is widely distributed in the nervous system as well as in many other peripheral organs, acting both as a neuromodulator and neuroendocrine releasing factor (Delgado et al., 2004; Iwasaki et al., 2019; Waschek 2013). In the brain, VIP regulates the secretion and expression of neurotrophic factors (Brenneman et al., 1990; Pellegri et al., 1998), has neuroprotective and anti-inflammatory effects (Delgado et al., 2004, Waschek 2013), and is up-regulated in neurons and immune cells after injury and/or inflammation (Armstrong *et al*., 2004, Waschek 2013).

Owing to VIP’s versatile and widespread effects, it is presently difficult to pinpoint its possible role in Betz cells without additional functional assessments. However, the particularly large size of Betz cells may imply unique metabolic/ homeostatic demands, for which VIP may contribute in terms of neuroprotective effects. Betz cells, with their large body size and long myelinated axons, are amongst the most vulnerable neurons (Mattson and Magnus 2006), which are affected in ageing and motor neuron disease (Cleveland et al., 2001, Scheibel et al., 1977, Udaka et al., 1986). High energy requirement, and reliance on long distance axonal transport along with a large cell surface area, increases exposure to undesirable factors (Morrison et al., 1998), likely rendering Betz cells in need of extra regulatory protection. In line with this hypothesis, it has been shown that VIP mRNA has been induced in facial motor neurons of mice following application of an inflammatory stimulus directly to the nerve (Armstrong *et al*., 2004). Moreover, the expression of PV in Betz cells (Szocsics et al., 2021, Preuss & Kaas 1996) may add another layer of protection through calcium regulation. Consistently, the proportion of PV-containing Betz cells decreases with age in humans (Szocsics et al., 2023), making them more susceptible to degeneration (Scheibel et al., 1977).

VIP induces concentration-dependent glycogenolysis in mouse cortical slices, which may indicate a possible regulatory mechanism for the local control of energy metabolism (Magistretti et al., 1981). The presence of VIP in the largest pyramidal cells may also have a relationship with the high demands of these cells for energy substrate. Such hypothesis is consistent with the maximal VIP staining in the largest pyramidal cells, which are located more medially in A4ab. Whether VIP being released from the Betz cells along with the other neurotransmitters or its functions are limited to the cytoplasm remains to be understood.

### 4.2. VIP+ interneurons in the motor cortex

VIP^+^ interneurons among the main subclasses of GABAergic neurons, being primarily located the supragranular layers of the cortex in the areas examined in this study, similar to observations in other species (Gabbott and Bacon 1997; Peters et al., 1987; Prönneke et al 2015; Tremblay et al., 2016, Kim et al.., 2017). VIP^+^ neuronal density across cortical layers showed a sharp decline toward the white matter, with much fewer VIP^+^ interneurons present in the infragranular layers. Supra- and infragranular VIP^+^ interneurons may have different functional roles as the two populations are transcriptionally distinct in mice (Wu et al., 2022).

Although the drop in the density of VIP^+^ inhibitory neurons in the infragranular layers was consistent in the three examined areas, the mean average density of VIP^+^ interneurons and their relative proportion to the total neuronal density were higher in the A4c compared to the areas A4a and A4b. This may reflect regional differences in network organisation for facial versus limb and body movements. Together with the differences in expression of VIP in Betz cells, this result further highlights the fact that functionally defined cortical areas are not necessarily homogeneous, a principle which has been demonstrated in terms of cellular structure (e.g. present results; Ungerleider and Desimone 1986), connections (e.g. Palmer and Rosa 2006; Rosa et al. 2009) and physiological properties (Yu et al. 2010; Yu and Rosa 2014).

Our density estimates for VIP^+^ interneurons in the motor cortex (3,065-4,745/mm^3^) are 50-100% higher than those in mouse motor cortex (Kim et al., 2017). Overall, we report a 5-7% contribution of VIP^+^ interneurons to the neuronal population in the motor cortex in the marmoset monkeys. Based on the 23% contribution of GABAergic interneurons in marmoset’s motor cortex (Bakken et al., 2021) and total neuronal density (Atapour et al., 2019), we estimate that VIP^+^ interneurons of the motor cortex constitute around 20-29% of the total interneuron population. These estimates are higher than those reported in several cortical areas of mice, which are around 11-15% of total GABAergic neurons (Gonchar et al., 2007, Pfeffer et al., 2013, Prönneke et al., 2015; Szadai et al., 2022). The observed morphologies of VIP^+^ interneurons in the marmoset motor cortex were similar to those reported in other species (Apicella et al., 2022; Gabbott and Bacon 1997, Markram et al., 2004).

Considering the soma size of neurons, the arbitrary fluorescence intensities in the VIP^+^ interneurons were similar to the Betz cells, indicating a comparatively similar expression of VIP. While the modes of connectivity of VIP^+^ interneurons with Betz cells remain to be elucidated, current literature suggest that VIP^+^ interneurons provide inhibitory inputs to pyramidal neurons mainly through forming disinhibitory circuits (Pfeffer et al., 2013; Pi et al., 2013, Lee et al., 2013). However direct inhibition of pyramidal neurons by this population has also been reported (Zhou et al., 2017; Garcia-Junco-Clemente et al. 2017; Yetman et al. 2019).

The abundance of studies on VIP^+^ interneurons in rodents has shed lights on the specific roles of this neuronal population, which include motor integration (Lee et al., 2013; Fu et al., 2014; Yu et al., 2019) and gain control (Pi et al., 2013). However, we are far from attaining a similar understanding about their function in primates, where only limited studies have been made available (e.g. Benson et al., 1991; Gabbott and Bacon 1997; Ong & Garey, 1991).

### 4.3. Conclusion

The distribution of VIP^+^ neurons in the marmoset motor cortex demonstrates that this neuropeptide is expressed both in interneurons and large pyramidal cells with putative projections to the spinal cord (Betz cells). The specific distribution of VIP in Betz cells hints at a different set of functions for this neuropeptide, as well as differences in neuronal circuitry involved in generating and controlling different types of body movements.

## ACKNOWLEDGEMENTS

Funding was provided by the National Health and Medical Research Council to Marcello G. P. Rosa (APP1194206) and Nafiseh Atapour (APP2019011). The authors acknowledge the contributions of the Monash Micro Imaging facility for providing training and support for confocal imaging and the Monash Histology platform for slide scanning services.

## CONFLIC OF INTEREST STATMENT

The authors have no conflicts of interest to declare.

